# Adult diet strongly affects cuticle thickness and its injury resistance in the insect *Narnia femorata* (Hemiptera: Coreidae)

**DOI:** 10.1101/2024.10.22.619666

**Authors:** EV(Ginny) Greenway, Tamsin Woodman, Zachary Emberts, Simon Chen, Walter Federle, Christine W. Miller

**Author notes:** Correspondence: Christine W. Miller.

## Abstract

Arthropods are the most diverse phylum on earth, accounting for up to 90% of animal species. The cuticular exoskeleton has played a vital role in their evolutionary success, but we know surprisingly little about the factors influencing its development, structure, and biomechanical properties. In this study, we examined whether and how nutrition affects the cuticle after an insect has completed its final molt into the adult stage. We found that a high-quality adult diet provided to the leaf-footed cactus bug, *Narnia femorata* (Hemiptera: Coreidae), over three weeks led to 4.1-times thicker cuticle with 3.7-times greater injury resistance relative to those that consumed a poor, but still ecologically relevant, diet. We further discovered that a high-quality adult diet allowed compensation for a sub-optimal juvenile diet. The cuticle of males was more injury resistant than that of females, as expected due to the aggressive male competition in this species. Altogether, our results show that, even after the final molt, nutrition can strongly influence the development and properties of the arthropod cuticle, a phenomenon that likely has drastic fitness-related consequences for locomotion, predator-prey interactions, and resource acquisition.

## Introduction

Arthropod cuticle is the second most common natural composite material in the world and provides one of the most evolutionarily successful forms of bodily protection. All parts of the exoskeletons of insects and other arthropods are built of cuticle, and it is involved in almost every biological function (1). Thus, it is surprising how little we currently understand about how cuticle properties are impacted by nutrition. A robust and impermeable cuticle is the first line of defense against the outside world, including biotic factors, such as competitors, predators, parasites, and pathogens, and abiotic stressors such as temperature and desiccation (2). The cuticular exoskeleton is also essential for stabilization of the body during terrestrial locomotion and flight, as well as for the functioning of the mechanical systems involved in movement, defense, reproduction, sensory perception, feeding and respiration (3). Further, thickening of the cuticle is one of the mechanisms by which arthropods evolve pesticide resistance (4).

After an insect molts into adulthood, the cuticle is typically hardened via a process called sclerotization (5). With this hardened “shell” on the exterior, the insect ceases external growth (1). From the outside, it appears that the insect’s growth and development are finished and that the sensitive period during which cuticle structure can be influenced by environmental factors (such as nutrition) is over (6, 7, but, see 8). Yet, careful histological studies have revealed that cuticular growth continues within the insect during early adulthood (9–11). Layer upon layer of procuticle are added internally, lining the body cavity of the insect and placing a limit on the extent cuticle can expand (11). Indeed, existing data suggests that 30 – 60% of the procuticle is produced after the adult molt in some species (12, 13). Insects also can modify the adult cuticle during the process of wound healing (14–16). Yet, it is largely unknown to what extent external factors influence the adult cuticle deposition process (but, see 17) and if there are functional consequences.

Our objective here was to examine the effects of the early adult diet on the thickness and injury resistance of insect cuticle. Our focal species was the leaf-footed cactus bug, *Narnia femorata* Stål (Hemiptera: Coreidae), an insect where a robust cuticle is likely to be especially valuable. Males in this species engage in vigorous physical contests over mating opportunities (18, 19). Cuticular injuries, including punctures and fractures, are common (Raina et al., in review; 20). We manipulated the nutrition of newly molted adult insects, reflecting natural dietary differences in the wild. We then examined the thickness, mass, and injury resistance of cuticle across the sexes and across three bodily regions. We examined both the outer epicuticle and underlying procuticle, which together comprise the insect exoskeleton. We predicted that insects allowed to feed on a high-quality adult diet for a longer time would develop a thicker procuticle with greater injury resistance, and that a high-quality adult diet would compensate for juvenile dietary deficiency. We predicted that males would be more injury resistant than females because of the prevalence of aggressive male-male competition in this species and that male hind legs would be the most injury resistant bodily region due to their role in physical contests. Finally, we tested whether the male hind leg cuticle was especially responsive to dietary quality, as is often seen in other attributes of sexually selected weapons and ornaments (21–23).

## Results

### A high-quality adult diet led to a thicker cuticle

We provided a sub-optimal diet to a focal group of juvenile *Narnia femorata*. Upon reaching adulthood, these insects were randomly assigned to one of three dietary regimes that varied in quality (Fig. 1). An additional, comparison group of insects were provided a high-quality diet throughout both juvenile and adult life stages so we could understand to what extent nutrition during early adulthood can compensate for earlier dietary deficiencies. All diets provided were ecologically relevant, mirroring natural, seasonal changes in their host plant, prickly-pear cactus.

**Fig. 1.**
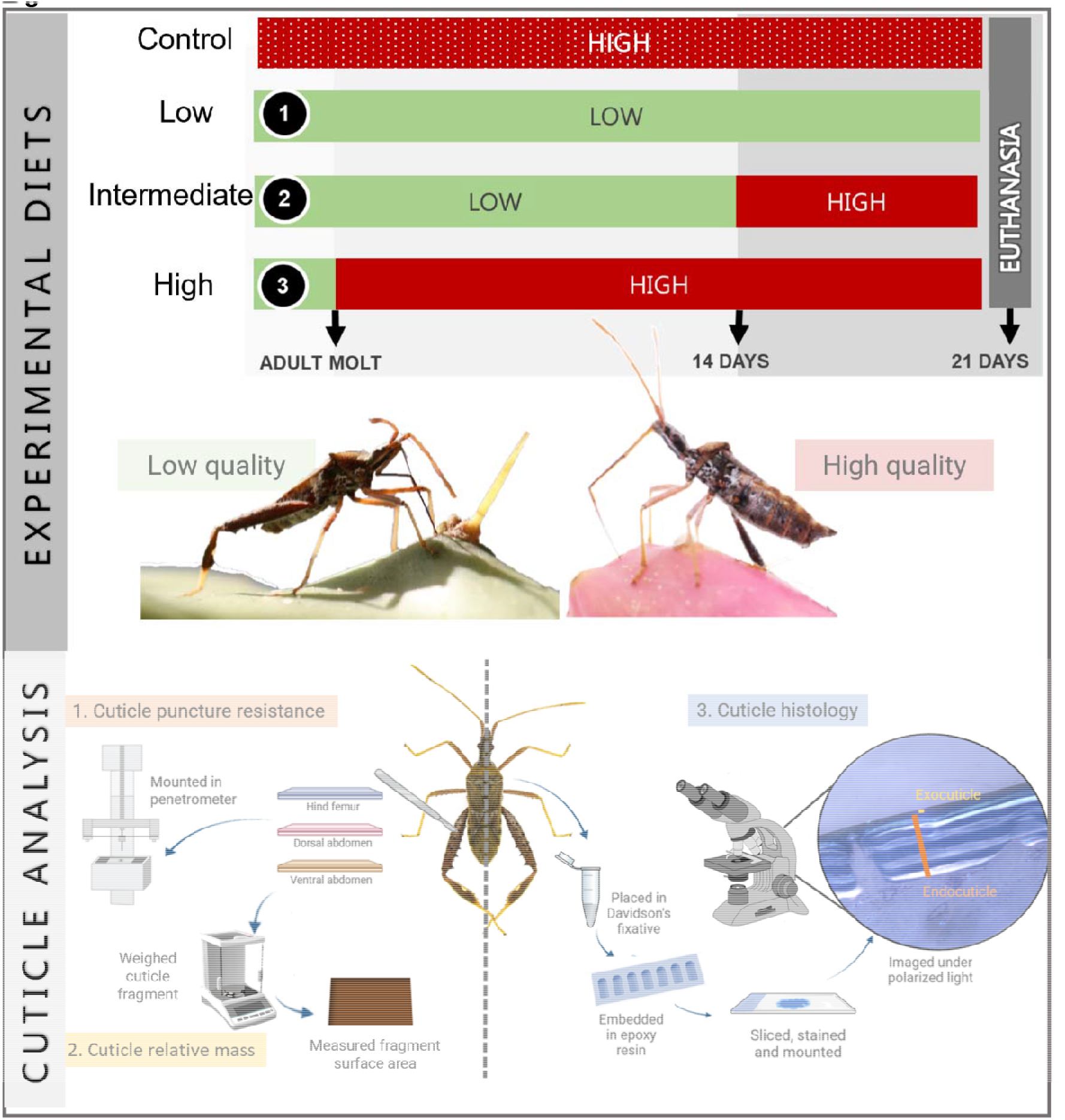
Diet quality manipulations (top) and subsequent cuticular measurements (below). Our focal insects were reared on suboptimal nutrition as juveniles (a mixed diet) then transferred onto a low-quality (1), an intermediate (2), or a high-quality (3) cactus diet regimen as adults. A subset of individuals (“Control”) was reared as a comparison group on consistent high-quality nutrition throughout juvenile and adult stages. We measured cuticle injury resistance via puncture tests, we harvested cuticle fragments and measured their surface area and mass, and we examined cuticle thickness via histology and microscopy.

We examined the insects’ exoskeleton 21 days after the adult molt, focusing on the procuticle, which is the bulk of the cuticle and includes both exocuticle and endocuticle. We also examined the epicuticle, the outer-most layer of the exoskeleton. We discovered that the procuticle thickness was dramatically impacted by our adult experimental dietary manipulation (Comparing low, intermediate, and high dietary treatments; n = 31; Linear Model (LM), Likelihood Ratio Test (henceforth, LRT), F_2,27_=20.9, P<0.001; Fig. 2); insects with access to the high-quality diet during adulthood had 4.7-times thicker procuticle relative to those provided the low-quality diet. We found no statistically significant difference in pronotum width between the three focal treatment groups, as expected (LM, F_2,63_ =0.87, P=0.426; Fig. S2). Higher-quality nutrition led to an increase in both the number of procuticle layers (Generalized LM, LRT, F_2,27_=5.25, P= 0.012; Fig. 2C) and their mean thickness (LM, LRT, F_2,27_=17.0, P<0.0001; Fig. 2D).

**Fig. 2.**
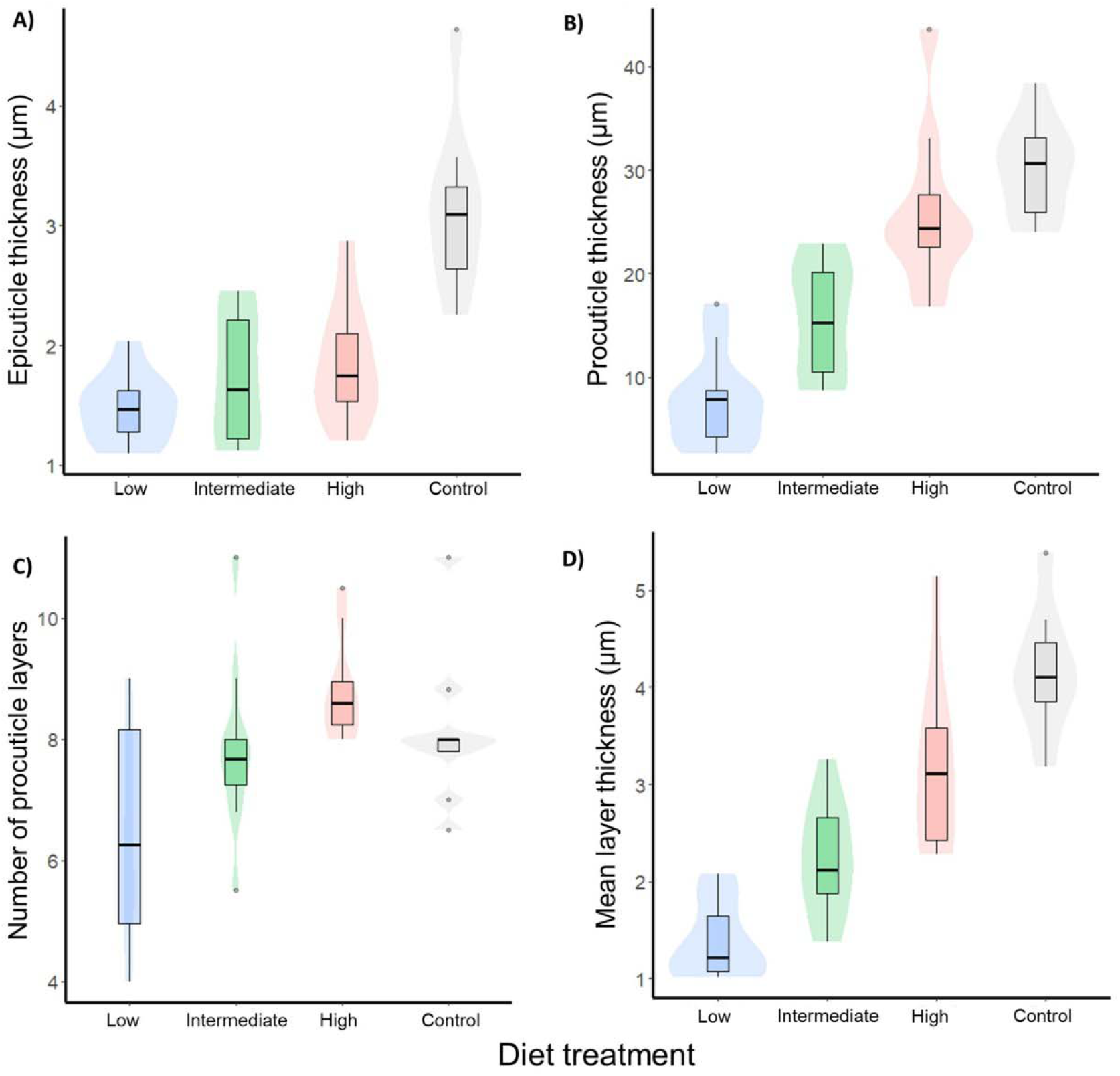
The impact of adult dietary regime on A) epicuticle thickness, B) procuticle thickness, C) procuticle layer number, D) mean procuticle layer thickness. Control individuals, those with a consistent high-quality diet throughout juvenile and adult life, are presented for qualitative comparison. Measurements of cuticle thickness were obtained from the ventral abdominal cuticle. Results shown are not adjusted for differences in body size; note that control individuals had larger adult body sizes (Fig. S2). Box plots show median and interquartile ranges, whiskers denote 1.5x interquartile range and points represent any data which falls outside of this range.

Additional findings from our three focal treatments showed that the thickness of the epicuticle and procuticle was positively associated with the body size achieved at the final adult molt (procuticle: LM, LRT, F_1,27_= 10.07, P=0.004; epicuticle: LM, LRT, F _1,27_ =43.79, P<0.001, Fig. 3). Males developed thicker epicuticle than females, but we did not find a sex difference in procuticle thickness (epicuticle: LM, LRT, F_1,27_ =11.92, P=0.002; procuticle: LRT, F_1,27_ =3.10, P=0.09; Fig. 3). Even though procuticle thickness was influenced by adult diet, we did not find evidence that adult diet influenced epicuticle thickness (LM, LRT, F_2,27_ =0.308, P=0.738; Fig. 2A, Fig. 3). Underlying raw data distributions are shown via background shaded violin plots.

**Fig. 3.**
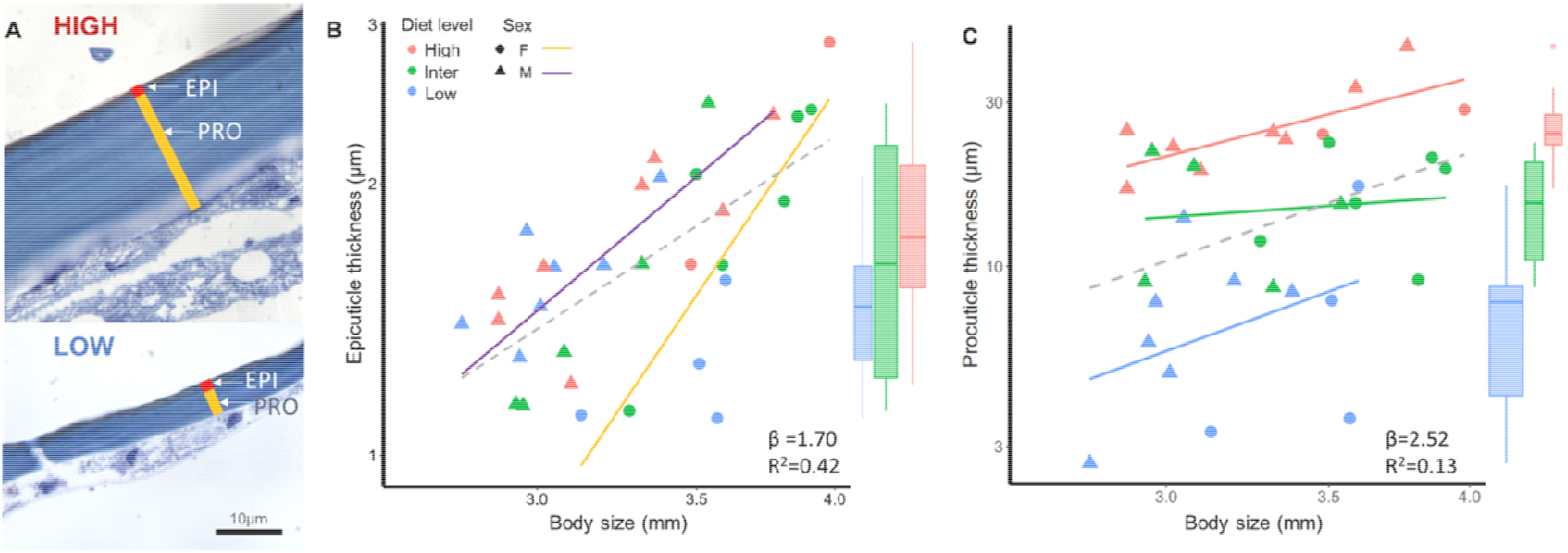
A) Ventral abdominal cuticle of *N. femorata* reared on a high-quality versus a low-quality diet (both x100 magnification). Epicuticle (EPI) thickness is indicated by the red bar and procuticle (PRO) is indicated by the yellow bar. **B)** Epicuticle thickness was greater for those at larger body sizes, but we did not find evidence it was affected by adult diet. **C)** Insects with a high-quality adult diet grew very thick procuticle for their body size, while those on the lower quality diets grew progressively thinner procuticle. In B) and C) β denotes the overall slope of the OLS regression (grey dashed line) for all points. These lines show the overall relationship between body size (measured as pronotum width) and cuticle thickness; the overall OLS slopes did not differ from isometry. R^2^ denotes goodness of fit of regression overall. Solid colored lines show OLS regressions for each sub-group.

Our next step was to examine the degree to which high-quality nutrition during adulthood allowed insects to make up for deficits in their earlier cuticle development. We compared the cuticle thickness of two groups, (1) insects reared on a sub-optimal diet as juveniles and placed onto a high-quality diet as adults (“#3 High” in Fig.1), and (2) insects with consistent access to high-quality nutrition (“Control” in Fig.1). It is well established that juvenile nutrition can have a pronounced effect on adult insect body size (24). Along these lines, we found that the control insects, with their higher-quality juvenile diet, grew larger bodies (35% larger pronotum width; Fig. S2) with 87% thicker cuticle on the ventral abdomen (Fig. 2). We discovered that a return to a high-quality diet in adulthood led to compensation in procuticle thickness for the juvenile dietary deficiency (with, as expected, no compensation of body size; Fig. S2); this effect was clear with and without correction for body size (Table S1; no statistically significant difference in the Control vs High-quality contrast).

### A high-quality adult diet improved injury resistance

We next assessed dietary effects on the resistance of the cuticle to injury caused by puncture. We dissected 2x3mm tissue-free cuticle sections from the hind femur, ventral abdomen, and dorsal abdomen of 66 experimental insects, including those tested above. Our custom-built penetrometer drove a sharp metal pin through each cuticle fragment while recording the force required to puncture the cuticle. We measured the surface area of the sections and calculated area-specific mass by dividing the air-dried mass of each fragment by its surface area. We found that puncture resistance varied over 180-fold across individuals, requiring from as little as 0.8mN up to 149.1mN of force for cuticle penetration across body parts. Insects with the high-quality adult diet developed on average 3.7-fold greater overall puncture resistance compared to those provided the low-quality diet (Linear Mixed Model (LMM), LRT, χ^2^= 39.8, P<0.001; Fig. 4).

**Fig. 4.**
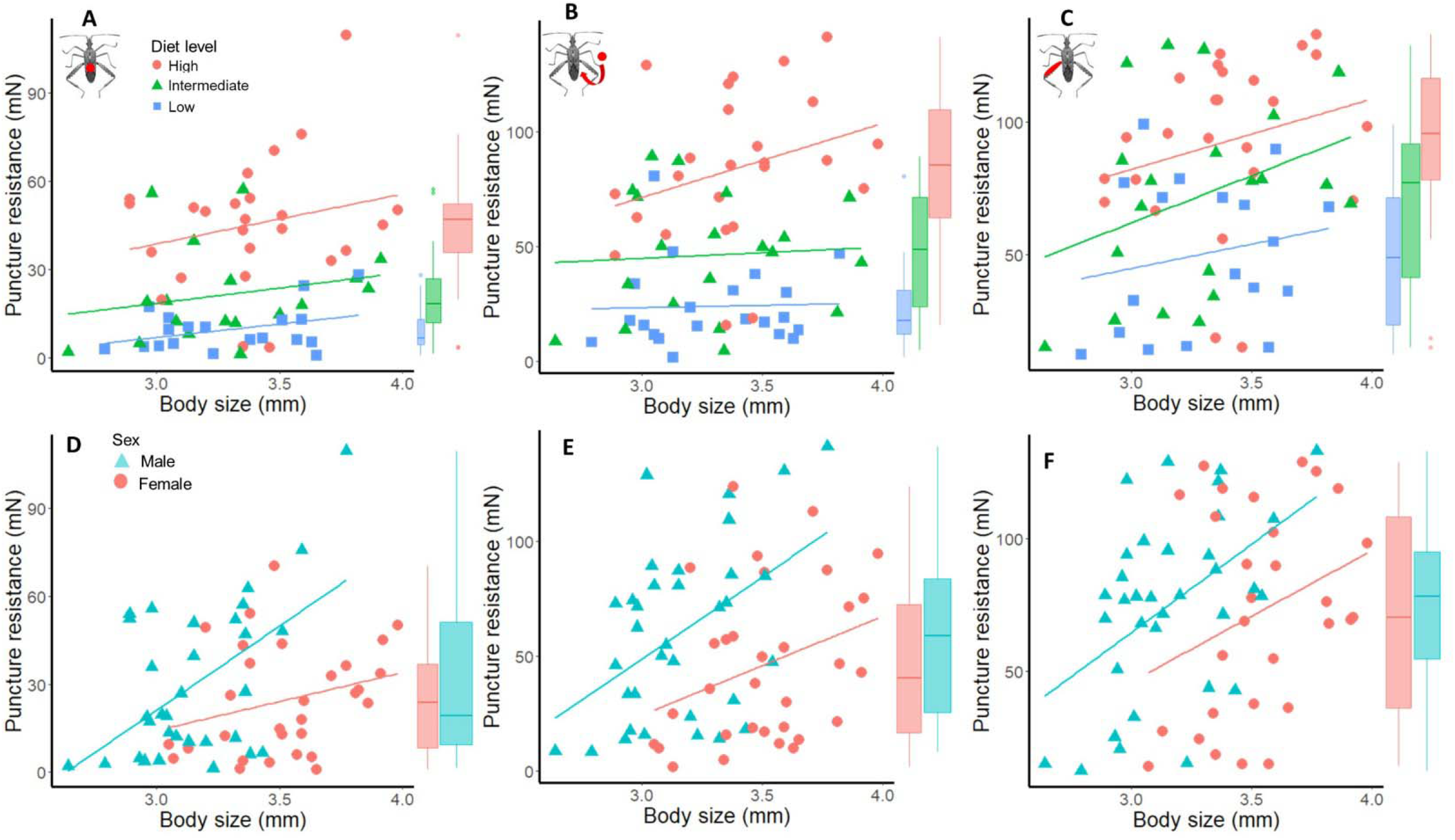
The impact of diet treatment on the relationship between cuticle puncture resistance and body size (pronotum width) across A) the dorsal abdomen, B) ventral abdomen and C) hind femur, alongside the influence of sex on puncture resistance across these same body parts (D, E, F). Separate regression lines are plotted for each diet and sex for visualization purposes.

Overall, larger individuals were more resistant to injury than small individuals across all body regions (LMM, LRT, X^2^ =14.6, P<0.001, Fig. 4). Within individuals, we found considerable differences in injury resistance based on the location from which cuticle was sampled. Hind leg cuticle was more puncture resistant than ventral abdominal cuticle, which in turn was more injury resistant than dorsal abdominal cuticle (Fig. 4). When adjusting for size differences between females and males (females are typically larger), we found that males, relative to females, showed greater injury resistance overall (LMM, LRT, X^2^ =12.26, P<0.001, Fig. 4) and across all three body parts (Table 1, Fig. 4).

**Table 1.** ANCOVA with Type II sums of squares testing the independent effects of body size (pronotum width), diet treatment and sex on the puncture resistance of the i) hind leg, ii) ventral abdomen and iii) dorsal abdomen cuticle (full model= Puncture resistance∼ Body size + Treatment + Sex).

| i) Hind leg |  |  |  |  |
| --- | --- | --- | --- | --- |
| *Factor* | *DF* | *Sum Sq* | *F-value* | *P* |
| **Body size** | 1 | 9287 | 10.24 | 0.00223 |
| **Diet treatment** | 2 | 15175 | 8.37 | 0.00064 |
| **Sex** | 1 | 5023 | 5.54 | 0.02199 |
| **Residuals** | 58 | 52580 |  |  |
| ii) Ventral abdomen |  |  |  |  |
| *Factor* | *DF* | *Sum Sq* | *F-value* | *P* |
| **Body size** | 1 | 6049 | 10.10 | 0.0023 |
| **Diet treatment** | 2 | 34702 | 28.96 | <0.0001 |
| **Sex** | 1 | 7489 | 12.50 | 0.0007 |
| **Residuals** | 61 | 36543 |  |  |
| iii) Dorsal abdomen |  |  |  |  |
| *Factor* | *DF* | *Sum Sq* | *F-value* | *P* |
| **Body size** | 1 | 2900.7 | 12.03 | 0.0010 |
| **Diet treatment** | 2 | 11959.9 | 24.79 | <0.0001 |
| **Sex** | 1 | 2465.6 | 10.22 | 0.00227 |
| **Residual** | 57 | 13750.1 |  |  |

Finally, we examined cuticle thickness, the area-specific mass of the cuticle, and puncture resistance in concert (Fig.1). As expected, cuticle thickness (obtained from histology) was closely related to the mass per unit area of cuticle taken from the same body region (Fig. 5A, log-log OLS regression, R^2^= 0.87, slope = 0.81, difference to isometry (β=1): P=0.002). We found that puncture resistance increased more than linearly with area-specific cuticle mass (Fig. 5, log-log OLS regression, R^2^= 0.81, slope = 1.38; difference to direct proportionality (β =1): P<0.001), indicating that thicker cuticle led to a more-than-linear increase in injury resistance Cuticle thickness and mass measurements were taken from histological sections of ventral abdominal cuticle. Cuticle mass was divided by the area of each cuticular section.

**Fig. 5.**
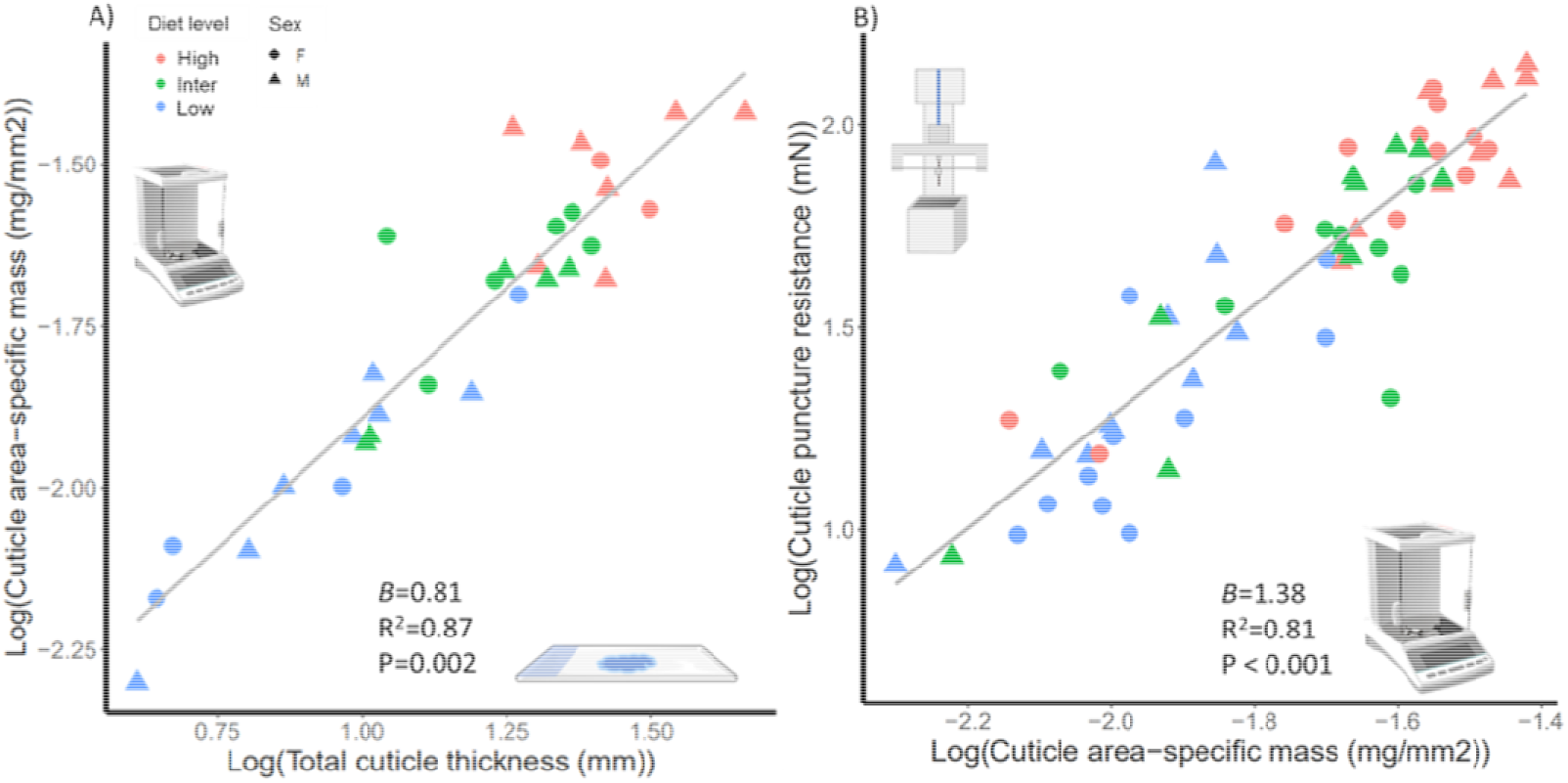
(A) Insects with thicker cuticle also had heavier cuticle, though the relationship scaled below isometry. (B) Insects with heavier cuticle had heightened resistance to puncture injury. β denotes the scaling exponent and P-value illustrates statistically significant deviation from isometry (β =1). One outlier (likely a measurement error) was excluded from B) when calculating slope values to optimize model fit. See Fig S3 for the equivalent figure containing the outlier.

### The abdominal cuticle, not the hind leg, was the most responsive to diet

The impact of diet on cuticle injury resistance differed across body regions (Treatment*Body region, LMM, LRT, X^2^ =14.40, P=0.006), suggesting variation in condition dependence across traits. To examine this further, we measured condition dependence as the relative difference in trait value between focal individuals given a high-quality adult diet versus those on the low-quality adult diet. Since the male hind leg is used as a sexually selected weapon in intrasexual contests, we predicted it would receive the greatest boost in injury resistance with improved nutrition. Yet, we found that the hind leg cuticle was the least condition dependent in both sexes (Fig. 6, Table 2). Whilst a high-quality diet caused an approximately 85% increase in injury resistance in both male and female hind leg cuticle, dorsal abdomen cuticle injury resistance increased by nearly 570% in males and 237% in females (Table 2). Surprisingly, we also did not see an overall effect of sex on condition dependence (Treatment*Sex interaction: LM, LRT, X^2^ =0.112, P= 0.737, Fig. 6). Overall, a high-quality diet resulted in consistently increased cuticle puncture resistance for the same body sizes, across all three body parts (Fig. S4).

**Figure 6.**
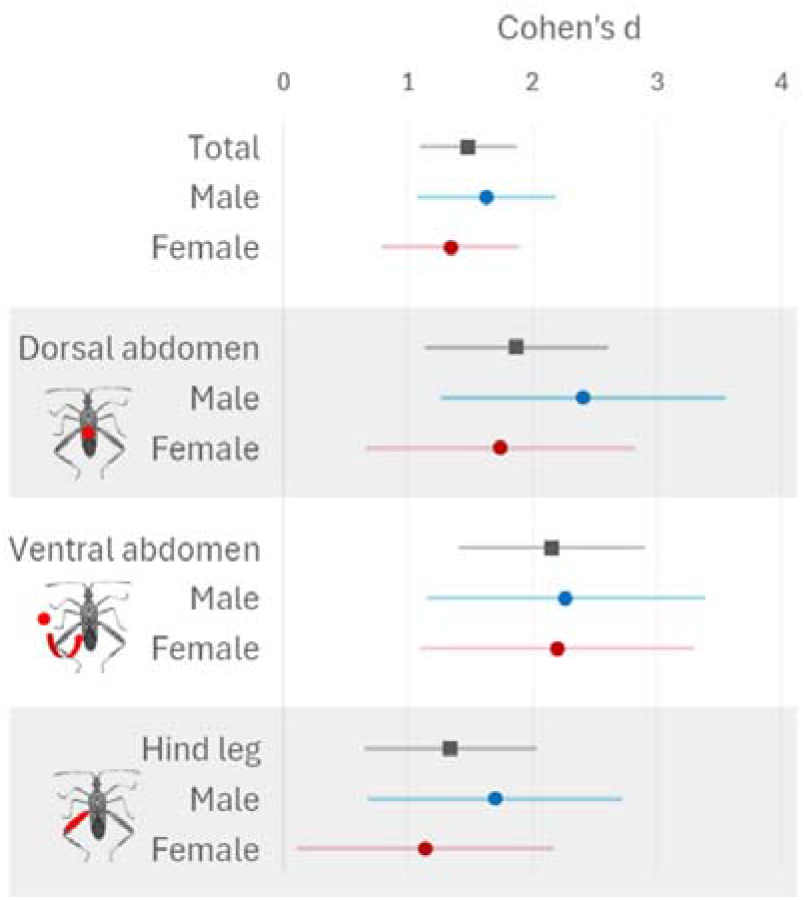
The magnitude of impact of diet quality (high versus low) on cuticle puncture resistance overall across all individuals and regions (total), by each sex, and then separately across the three body parts tested. Points denote Cohen’s d effect sizes with associated 95% confidence intervals.

**Table 2.** To examine the relative condition-dependence of traits, we examined changes in trait values between low- and high-quality adult diets. Results are presented as raw differences in values, standard errors and the associated percent increase for those fed the high-quality diet.

|  | Males |  |  |  |  | Females |  |  |  |  | Diet * Sex |
| --- | --- | --- | --- | --- | --- | --- | --- | --- | --- | --- | --- |
|  | Raw difference | % change | SE | *t* | *P* | Raw difference | % change | SE | *t* | *P* | *F* and *P*-value (*d.f.*) |
| Hind leg puncture resistance (mN) | **44.2** | **+84.9** | **10.94** | **-4.037** | **0.0001** | **40.4** | **+84.2** | **16.18** | **-2.497** | **0.0224** | 0.018, 0.894 (1,38) |
| Dorsal abdomen puncture resistance (mN) | **43.4** | **+569.7** | **7.58** | **-5.723** | **<0.0001** | **27.5** | **+236.7** | **6.98** | **-3.939** | **0.0009** | 1.86, 0.18 (1,39) |
| Ventral abdomen puncture resistance (mN) | **62.4** | **+215.1** | **11.58** | **-5.388** | **<0.0001** | **57.0** | **+301.2** | **10.83** | **-5.264** | **<0.0001** | 0.032, 0.859 (1,41) |
| Ventral abdomen area-specific mass (mg/mm<sup>2</sup>) | **0.0199** | **+180.9** | **0.002** | **-8.713** | **<0.0001** | **0.0128** | **+112.3** | **0.003** | **-4.393** | **0.0003** | 3.55, 0.067 (1,37) |
| Body size (pronotum width, mm) | 0.1406 | +4.5 | 0.104 | -1.358 | 0.189 | 0.0781 | +2.25 | 0.106 | -0.739 | 0.468 | - |

Statistically significant differences in trait values between individuals on low and high-quality dietary regimes are highlighted in bold. Within each sex, traits revealed positive condition dependence, but males and females did not respond differently, as shown by no significant diet by sex interaction terms. The puncture resistance of the male dorsal abdomen showed the greatest increase with a high-quality diet.

## Discussion

We found that a high-quality diet during the first three weeks of adulthood resulted in a thicker, more injury resistant cuticle in *Narnia femorata.* Young adult insects with limited access to high-quality nutrition, as could occur, for example, seasonally or with anthropogenic habitat degradation, may have increased susceptibility to injuries from predators, parasites, competitors, and disease (13, 25, 26). Furthermore, a thin, weak cuticle may not provide the musculoskeletal support necessary for efficient locomotion and flight (27, 28), potentially limiting dispersal and gene flow. Dietary effects on the insect cuticle, if common, may have vast consequences for life-histories, evolutionary trajectories, and population persistence. Yet, these effects and their consequences are only just beginning to be examined (6, 8).

Very little is known about the causes and consequences of within-species differences in the arthropod cuticle, even though detailed investigations of the cuticle were well underway by the middle of the 20^th^ century (e.g., 12, 29). During this time, Neville (9–11) revealed that the outside appearances of insects can be deceiving; the cuticle keeps growing layer-by-layer on the inside even during adulthood. We have since learned that procuticle thickness can increase rapidly, by up to 25 times, during the first few weeks of adulthood (13, 30). The cuticle is dynamic in other ways during early adulthood; for example, cuticular injuries can be repaired (16, 31). The deposition of procuticle layers can be dependent upon whether or not food is available and the presence of specific key compounds (32–35). Along these lines, a number of insect species rely on symbionts for biosynthesis of compounds required to thicken and harden cuticle (13, 36).

Recent work has shown that elevated ocean temperatures can drive short-term effects on arthropod cuticle hardness and stiffness (37). Yet, work on the influence of environmental factors on cuticle formation and material properties has largely stopped there (38).

Environments in nature are rarely static, and fluctuations in resource quality and availability are common. In the designing of this experiment, we prioritized treatments with ecological relevance.

*N. femorata* encounters dynamic plant diets naturally in the wild as the seasons change. It feeds on the reproductive structures of *Opuntia* cacti, passing multiple, overlapping generations as cactus fruit is produced in the spring and until it is red and ripe in the autumn. Thus, each generation hatches to find cactus fruit in a different phenological stage. Previous work has shown that feeding on this dynamic resource leads to numerous phenotypic effects on morphology and behavior (39–41). Further, Woodman et al (34) showed that an improved diet during the juvenile stages can lead to a 4.4-times increase in the injury resistance of the adult cuticle. In the current study, we asked if *adult* diet (not only juvenile diet) matters to the development of the insect cuticle and its properties. Our diets for this study were centered around the presence and absence of red, ripe cactus fruit (Fig. 1). *N. femorata* can easily become stranded without this high-quality food late in the season because deer, tortoises, birds, and rodents also prize red, ripe cactus fruit (40). Relative to Woodman et al. (34), this study took an expanded view, examining multiple body regions across both sexes and measuring cuticular layers using histology. We further tested the extent to which a return to good nutrition in adulthood results in compensatory growth of the cuticle.

### A high-quality adult diet led to a thicker cuticle

Many insects lay down layers of cuticle for the first weeks of adulthood (11). In fact, counting cuticle layers is one means to estimate the age of individual insects (42, 43). Here, we found that a high-quality adult diet led to a 4.1-times thicker procuticle by the third week of adulthood relative to a low-quality diet (Figs. 2, 3). We discovered that the increase in overall procuticle thickness was due to an increase in the thickness of individual layers as well as an increase in layer number (Fig. 2C & D).

We found evidence for compensatory growth in the cuticle during the adult stage. Our focal insects, those in the “Low”, “Intermediate”, and “High” treatments, experienced the same regime of sub-optimal diet as juveniles (Fig.1). Although they were largely stunted in overall body size from the suboptimal juvenile nutrition, the individuals moved onto the high-quality diet upon reaching adulthood were able to rapidly lay down procuticle. Indeed, we found that these experimental individuals on high quality adult diets were able to catch up in cuticle thickness to their counterparts who were consistently well-nourished throughout the juvenile and adult stages (Fig. 2B&C; Table S1).

We next examined the relationship between the thickness of cuticle, the area-specific mass of the cuticle, and its puncture resistance. We found them to be positively related (Fig. 5). As expected, cuticle thickness was closely related to the mass per unit area of cuticle taken from the same body region (Fig. 5A). Interestingly, the slope was less than isometry. Such a pattern could emerge if thicker cuticle is less dense than thin cuticle. Indeed, this could be based on the relatively larger proportion of endocuticle, which is typically less dense than the other components of the exoskeleton (44, 45). Cuticle with greater mass had greater resistance to injury (Fig. 5B).

### A high-quality adult diet improved injury resistance

Layers of insect procuticle include the nitrogen-containing polysaccharide, chitin, which, together with specialized, nitrogen-rich proteins, make the cuticle especially resistant to injuries, including those caused by puncture (46). We found that adult insects with greater exposure to a high-quality diet showed 3.7-times greater resistance to puncture injury (Fig. 5). We predicted that the hind leg cuticle would be the most injury resistant due to its central role in male-male sexual contests, both in offense and in defense (18, 19, 47). We indeed found that hind leg cuticle was more injury resistant than ventral abdominal cuticle, which in turn was more injury resistant than dorsal abdominal cuticle.

*N. femorata* exhibit a wide range of injuries after living in mixed-sex groups, a likely result of male aggression that targets both females and other males (20, Raina et al. in review). Forceful hind- leg squeezes are a characteristic component of male-male competition in this family of insects (18). Females are much less likely to initiate and engage in physical contests, though they can be a target of male aggression (20). Here, we found that, for a given body size, males were more injury resistant than females across all the body regions considered (Fig. 4, Table 1).

We were surprised to find such a pronounced difference in injury resistance across individuals. Injury was induced with as little as 0.8mN, while other specimens required up to 149.1mN, a shocking 177-fold difference. This difference likely has pronounced biological relevance. O’Brien and Boisseau (48) documented that male *N. femorata* can exert up to 50mN of hind-leg squeezing force during male-male contests, thus structures with injury resistance values below 50mN could be harmed in male-male contests. Indeed, the line of investigation described here was instigated because differences among individuals in cuticle stiffness were detected with just a slight squeeze of the human hand. The fact that some three-week-old adults have thin, pliable, and almost translucent cuticle evokes the vulnerable period that insects generally experience immediately after molting (49, 50). While a few days of vulnerability comes with challenges, this longer period of thin, pliable cuticle brings into question how those affected can walk, fly, mate, feed, and survive predation attempts. Data documenting intraspecific variation in insect cuticle thickness is scarce, but some species of polymorphic ants exhibit considerable levels of variance (ten-fold difference in cuticle thickness) between the smallest workers and large soldiers (51).

However, thin-cuticled ants, as well as thin-cuticled termites (52), are buffered from predation, desiccation and disease by the safety of the nest and their social environment, unlike free-living *N. femorata*.

### The male dorsal abdomen, not the hind leg, was the most responsive to diet

Life history theory predicts that the ornaments and weapons of sexual selection, as well as any other trait under directional selection, should evolve heightened condition dependence (53, 54). In other words, the degree of their expression should be especially sensitive to nutrition and other factors that impact physical condition. Male *N. femorata* often raise and hold their hind legs in the air before and during battles; thus, these limbs appear to have a signaling function. As expected, the size of these traits shows heightened condition dependence relative to other traits, including the homologous traits of females (41, 55). Here, we examined the injury resistance of the hind legs, specifically testing the degree to which this injury resistance responded to improved diet. We estimated relative condition dependence as the proportion change from the low-quality diet to the high-quality diet (23). Although males had greater injury resistance than females, we did not find evidence that the overall condition dependence of male traits exceeded that of female traits. (Table 2). The injury resistance of the dorsal abdomen of males was highly responsive to diet (+570%), much more so than their hind femur (+85%). These results are in line with others that suggest that heightened condition dependence is not ubiquitous in the weapons and ornaments of sexual selection (56). Increased condition dependence might be more commonly found in trait components that easily convey information to conspecifics (57), such as size, shape, and behavior. It is also worth exploring whether traits especially susceptible to attack and injury may also show a heightened response to improved physical condition. Intriguingly, the dorsal abdomen, covered with wings, is frequently a target of male-male competition in this family of insects (18) and can be commonly injured via puncture wounds from the sharp hind-leg spines of opponents (58).

### Nutritional constraints on insect cuticle formation

Many organisms encounter natural variation in their diet over time and space, and the results presented here show that such variation can have important implications for how insect bodies are built, with a myriad of possible fitness consequences. While we found here that the formation of cuticle layers, and the thickness of those layers, are influenced by diet, there is much that is not known. We do not yet know how widespread this phenomenon is across the insects, whether all insects stop laying down additional layers at a certain time point, and to what extent the timing of cuticle formation and the thickness of the layers is influenced by diet. Further, this work did not examine the precise nutrients required to grow a robust cuticle. The insect cuticle is rich in nitrogenous compounds (11), and the differences in cuticle seen here may be due to differences in the availability of nitrogen via amino acids and phenols (59–62). Indeed, Bernays (63) suggested that insects might conserve amino acids by investing less in their cuticle when needed. Herbivorous insects often receive inadequate amino acids or insufficient quantities of nitrogen from their host plants, and nutrient deprivation can become worse due to seasonal changes (64, 65). Future work is needed to manipulate specific dietary nutrients, including amino acid and phenolics, to determine precisely what is required for the building of a thick, injury-resistant cuticle.

### Conclusions

An enhanced focus on nutritional effects on arthropod cuticle is long overdue. This study has shown for the first time that the diet of an adult insect can have drastic effects on cuticular thickness and its injury resistance. The consequences of these findings may be profound, especially if representative of arthropods as a whole (66). While most animals on this planet have a skeleton made of cuticle, surprisingly little is known about how environmental factors influence its growth and development (38).The findings here beg the question – if diet affects the cuticle so strongly, what other environmental factors also have an effect?

## Materials and Methods

### Insect husbandry

Insects used in this study were the offspring of 10 male and 10 female *Narnia femorata* Stål, 1892 (Hemiptera: Coreidae) collected from the wild near Starke, Florida in August 2019. All cactus pads and fruit were harvested from the same North Central Florida location. We housed juveniles in groups of 2 to 17 individuals on a potted *Opuntia mesacantha ssp. lata* cactus cladode with red, ripe fruit. Upon reaching the penultimate (4th) instar, we transferred the 85 juveniles into their own container containing a potted cactus pad without cactus fruit. The loss of cactus fruit reflects what commonly occurs naturally when competitive herbivores (e.g., deer, tortoise, rodents, and birds) remove fruit from a cactus patch. Thus, our focal insects experienced a period of high-quality nutrition followed by low-quality nutrition as juveniles (hereafter, “sub- optimal” diet). One of our objectives was to understand the degree of compensation that can take place in early adulthood to make up for a reduced-quality juvenile diet. Thus, as a comparison group, we raised a subset of insects on a high-quality diet throughout both juvenile and adult life stages (hereafter “Control” diet).

When focal insects molted into adults, they were randomly assigned to one of three dietary regimes: 1) a high-quality diet for life, 2) a low-quality diet for life, 3) a low-quality diet for the first fourteen days of adulthood with a switch to the high-quality diet for the final seven days (Fig. 1). Twenty-one days after the molt to adulthood, insect pronotum width was measured as a proxy for body size (a representative measure of overall body size in this species; (39–41)) using digital calipers, then insects were cold-euthanized at -20°C. We then bisected insects along the midline of the body using a scalpel; one half of the body was returned to -20°C. Specimens were frozen for up to five months prior to testing. We submerged the other half of the body in Davidson’s fixative for ∼14 hours upon which time it was transferred to 70% ethanol and frozen following (38).

### Cuticle thickness

The fixed halves of each experimental insect were gradually transitioned from 30% to 100% acetone through intermediate concentration steps. Then, ventral abdominal samples from a standardized location were embedded into epoxy blocks using the Sigma-Aldrich Epoxy Embedding Medium Kit 45359 and standard protocol (following 67). Hardened epoxy blocks were sliced into 0.5-micron thick cross sections using a Leica EM UC7 Ultramicrotome, mounted on glass microscope slides and then stained using 0.1% methylene blue. We viewed and photographed (camera: Point Grey Grasshopper GS3-U3-51S5C-C) these cuticle cross sections under a Leica DMR-HC upright microscope using crossed polarizers and a 100x/1.25 oil objective to better resolve the layers of epicuticle and procuticle. We found that the cuticle thickness was somewhat variable within individuals. Thus, we estimated the mean thickness of each individual’s epicuticle and procuticle using photographs of the sections and Image J v1.52t (68). We then extracted the number and thickness of individual procuticle layers using a custom-written routine in MATLAB R2022b (Mathworks, Natick, MA).

### Injury resistance and cuticle area-specific mass

We measured injury resistance and the force required to puncture the cuticle using a 10 micron T20-100 Tungsten Probe Tip following (34), and using the same penetrometer. We selected this probe because of its tip radius of curvature is in the same range as that of the major leg spines in males (T20-100 tungsten probe: 3.265 +/- 0.312 µm (n=7), male spines: 2.472 +/- 0.803 µm (n=16)). When males in this species fight (19), they often press such leg spines into the ventral abdomen and other bodily regions of their opponents. A cuticle section of each frozen hind leg, ventral abdomen, and dorsal abdomen of approximately ∼2x3mm was removed from the same bodily location using a scalpel blade. These flat cuticle sections were clamped into an aluminum sample holder with a circular 1.2mm diameter hole through which a standardized location of the cuticle sample was exposed. With the penetrometer, the tungsten probe mounted on a steel bending beam was moved at a constant speed into the cuticle fragment. A small piece of reflective foil glued on the reverse side of the bending beam served as a target for a D12 fiber optic sensor mounted above the beam (Philtec, INC., Annapolis, USA). The sensor measures the distance between its tip and the reflective foil (and thus the deflection of the bending beam) as a voltage signal. The difference between the voltage at maximum deflection and the baseline voltage was used to calculate the puncture force for each cuticle sample. Samples were then stored frozen before being used for measurements of area-specific cuticle mass. Freezing cuticle samples did not impact their mass or mechanical properties (Supplementary methods; Fig. S1), consistent with Aberle et al. (69).

Attached fat bodies and the epidermis were removed from the ventral abdomen sections using forceps under a Leica MZ16 dissecting microscope. We obtained air-dried mass measurements using a MC5 microbalance (Sartorius, Göttingen, Germany) then transferred the sections to a compound microscope (Leica DMR-HC) to obtain images at 5x magnification and ultimately surface area measurements using ImageJ (68). We then calculated the area-specific mass of the ventral abdominal sections by dividing the mass of each fragment by its surface area.

### Statistical analyses

We first examined the impact of diet on ventral abdomen epicuticle and procuticle thickness, using linear models with the cuticle element of interest as the dependent variable and diet treatment, body size and sex as independent variables in each model (*lme4* package) and examined the significance of each term independently using the *drop1* function (70). We then took a closer look at individual layer thickness using a linear mixed model, with individual ID included as a random effect to account for multiple layer measurements per individual. To establish if diet impacted the number of procuticle layers individuals grew, we ran a linear model with average procuticle layer count (taken from between 2 to 8 measurements per individual) as the dependent variable. We then formally assessed how cuticle thickness scaled with body size using OLS regression, log-transforming both variables and testing whether the slope value differed from 1 (isometry) using the *car* package *linear-Hypothesis* function (71).

To ascertain whether individuals could compensate for sub-optimal juvenile nutrition once placed on a high-quality adult diet, we then repeated the same analysis but incorporated Control individuals to provide an ecologically realistic comparison. These Control individuals were provided with cactus fruit throughout their juvenile development (Fig. 1) and grew substantially larger than the focal experimental individuals across the three treatments (Fig. S1; F_3,81_=72.6, P<0.0001, Fig. S2), as expected (41). We ran linear models with and without body size accounted for as an independent variable, effectively comparing relative and absolute cuticle thickness across diet treatments, respectively. We again used linear regression models to assess the correlation between cuticle thickness and area-specific cuticle mass.

To capture functional effects of diet-induced changes in cuticle thickness, we then analyzed the impacts of sex, body size and adult diet treatment on cuticle puncture resistance across three different body parts, the ventral abdomen, dorsal abdomen and hind femur. We used the same linear model structure as above, examining the significance of each term independently using the *drop1* function. Interaction terms were non-significant and therefore removed from the model for parsimony. Finally, to examine the degree of condition dependence across different body parts, we focused on the differences in injury resistance between individuals raised on high-quality adult diets and low-quality adult diets (following approach in (23)). We re-ran a linear mixed model on this subset of data, incorporating a diet x body region interaction term and including individual ID as a random effect to account for the inclusion of multiple measures from each specimen. We then calculated the mean difference in puncture resistance between low- and high-quality diet individuals using the *psych* package summary function and determined what this difference was as a percentage of the low-quality diet trait mean values (72). We quantified variation in the impact of these two diets across body parts and sexes using Cohen’s D effect size coefficient, before formally testing for sex-specific condition dependence across traits by incorporating Sex x Diet interaction terms alongside body size in separate linear models for each body area. We then used the same OLS approach detailed above to examine if the log-log scaling relationship between puncture resistance and body size varied between individuals on low- and high-quality diets across the three different body parts. All statistical analyses were performed in R v4.3.0.

## Acknowledgments

This manuscript is based upon work supported by the National Science Foundation under Grant No. IOS-2226881. The authors thank the University of Manchester for the Industrial/Professional Placement Scheme that enabled T. Woodman to undertake insect rearing and data collection.

The authors also thank Dr Karin Müller at the Cambridge Advanced Imaging Center for guidance and assistance with histology.

## Author Contributions

Funding was acquired by C.W.M., and the study was conceived by C.W.M., Z.E., W.F., and S.C. S.C. built and maintained the penetrometer. T.W., E.V.G., and W.F. performed data collection. T.W. contributed a written report of preliminary findings, E.V.G. conducted statistical analyses and prepared the figures. The manuscript was written by E.V.G. & C.W.M. All co-authors helped improve the paper via reviewing and editing. **Competing Interest Statement:** The authors declare no competing interests

**This PDF file includes** the following: Main Text, Figures 1 to 5, Tables 1 to 2, Supplementary Fig. S1 to S4, Supplementary Table S1.

