## Supplementary Materials for "Adult diet strongly affects cuticle thickness and its injury resistance in the insect *Narnia femorata* (Hemiptera: Coreidae)"

**Impact of freezing on cuticle mechanical properties**

**Methods** We assessed the impact of freezing on measurements of injury resistance to better understand the biological relevance of the values we generated. Adults of *Leptoglossus phyllopus* (Hemiptera: Coreidae) were selected for this assessment because they are closely related to *Narnia femorata* and because live specimens were available at the time of biomechanical testing. Insects were reared in a mesh bug dorm (30cm^3^) and provisioned with *ad libitum* organic green beans and sunflower seeds and water-soaked cotton wool for moisture, under a 12∶12h day-night cycle at approximately 25°C and 40–50% humidity. Ten mature *L. phyllopus* adults were randomly selected for cuticle sampling. To establish the mechanical properties of live insects, individuals were cold anesthetized for 5 minutes (not frozen) and then euthanized by decapitatation. Using a scalpel, we immediately removed a fragment of cuticle approximately 2x3mm in size from the ventral abdomen, mounted it in the penetrometer and recorded its puncture resistance before measuring its surface area and mass. We then froze the remaining part of these individuals at -20°C for 7 days, before cutting another equivalent section of ventral abdomen cuticle and repeating the same measurements. This enabled us to compare the mechanical properties of the same insect’s cuticle with and without freezing. We found no evidence of a change in either relative mass (paired T test, *t* = -1.4481, df = 9, *P*= 0.1815) or puncture resistance (paired T test, t = 1.3882, df = 9, *P* = 0.1985) due to freezing. We further examined the puncture resistance and cuticle area-specific mass for four adult *L. phyllopus* that had been frozen for over 12 months. We included these samples to establish if cuticle properties changed with longer periods of freezing. This longer duration of freezing exceeded the period of freezing for *N. femorata* samples in our main study. We did not find evidence that cuticle properties changed over this longer time frame (Fig S1).

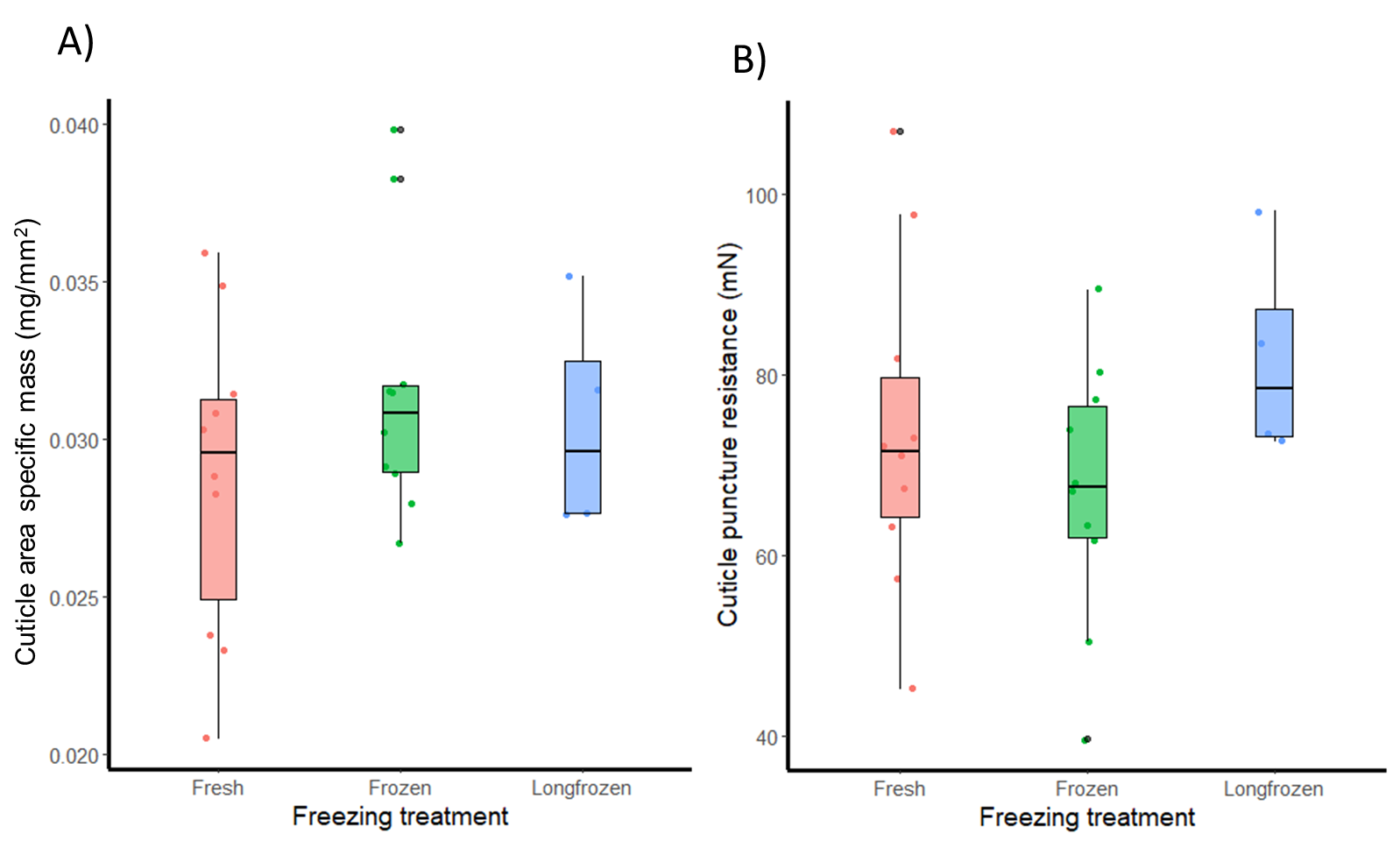

**Fig. S1.** No evidence that freezing for either 7 days (green) or upwards of a year (blue) impacts cuticle area-standardized mass (A) or puncture resistance (B) when compared to fresh cuticle samples (red).

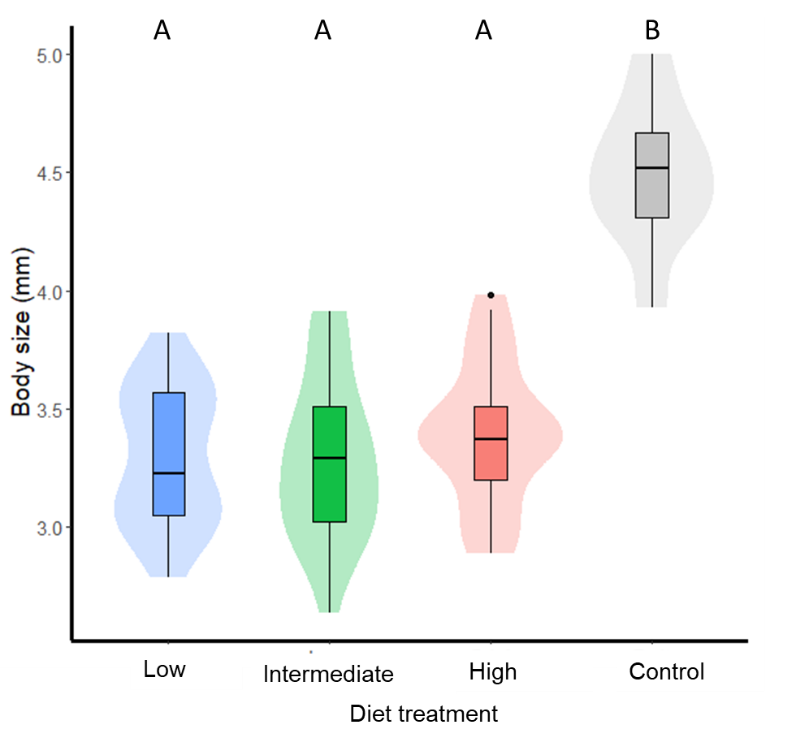

**Fig. S2**. Variation in body size (pronotum width) across the three experimental diet treatments compared to the control group which received access to ripe cactus fruit throughout development. Letters denote significant differences at the p<0.001 level. Box plots show median and interquartile ranges, whiskers denote 1.5x interquartile range and points represent any data which falls outside of this range. Underlying raw data distributions are shown via background shaded violin plots.

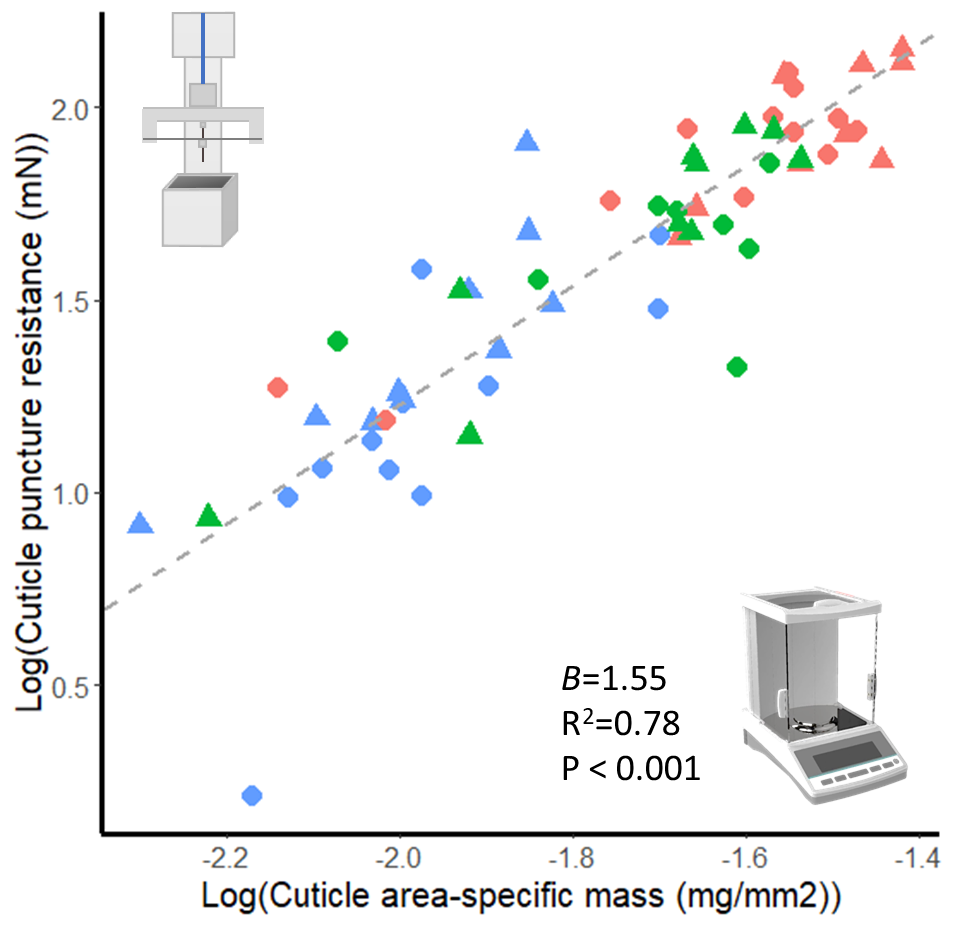

**Fig. S3.** Scaling relationship between ventral abdomen cuticle resistance and area-specific mass including outlying individual in lower left.

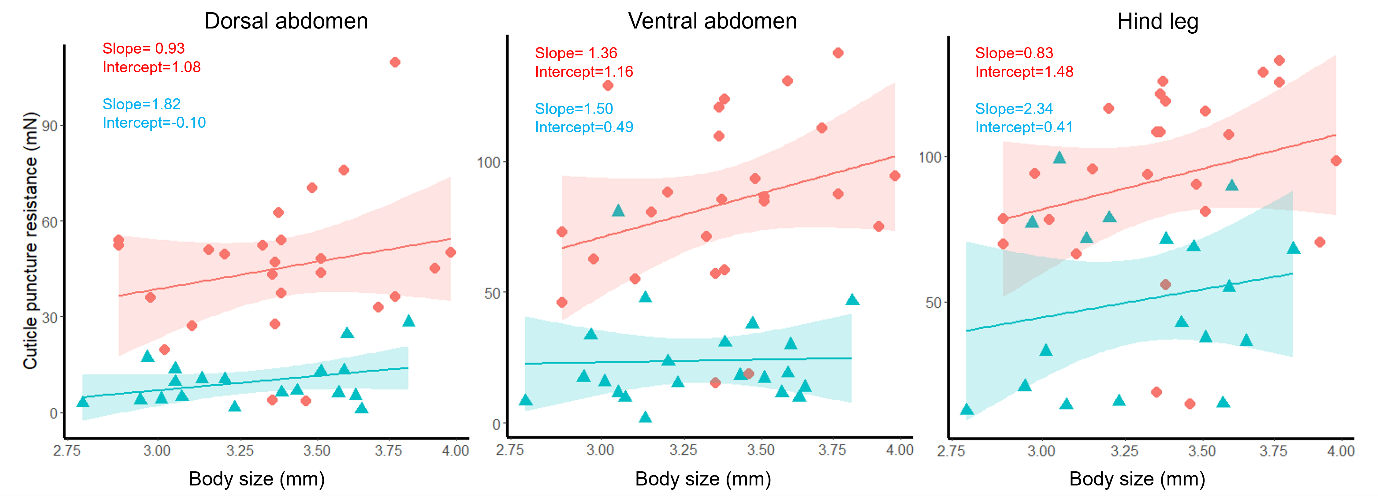

**Fig. S4.** The relationship between male body size (pronotum width) and cuticle injury resistance on high (red) and low (blue) diets across body parts. None of the OLS regression slope values differed significantly from isometry (slope=1) and there were no significant body size (pronotum width) x diet treatment interactions to report.

**Table S1.** We did not find evidence that experimental individuals feeding upon high-quality adult diets (but sub-optimal juvenile diets) differed in procuticle thickness relative to Control individuals raised on a consistent high-quality diet (Control-High contrast for procuticle). These results suggest that a high-quality adult diet can allow for compensation for a sub-optimal juvenile diet. These results also suggest that two weeks of high-quality nutrition (followed by one week of low-quality nutrition, altogether our “intermediate” diet regime) was not enough to compensate for the sub-optimal juvenile diet. Tukey posthoc contrasts of epicuticle and procuticle thickness across diet treatments, from Type II ANOVA models with (adjusted) and without (raw) body size included as an explanatory variable. Statistically significant differences between treatments are highlighted in bold. Single-step adjusted P-values reported.

| Pairwise comparison | Raw data | | Adjusting for body size differences | |
| --- | --- | --- | --- | --- |
| *Epicuticle* | t-value | P value | t-value | P value |
| Intermediate-Low | 1.123 | 0.678 | 0.288 | 0.991 |
| High-Low | 1.757 | 0.310 | 0.759 | 0.864 |
| Control-Low | **7.193** | **<0.001** | 0.701 | 0.889 |
| High-Intermediate | 0.642 | 0.918 | 0.490 | 0.958 |
| Control-Intermediate | **6.044** | **<0.001** | 0.617 | 0.921 |
| Control-High | **5.329** | **<0.001** | 0.349 | 0.984 |
| *Procuticle* |  |  |  |  |
| Intermediate-Low | **3.222** | **0.014** | **2.770** | **0.039** |
| High-Low | **7.285** | **<0.001** | **7.134** | **<0.001** |
| Control-Low | **8.750** | **<0.001** | 1.773 | 0.287 |
| High-Intermediate | **4.027** | **0.001** | **4.549** | **<0.001** |
| Control-Intermediate | **5.570** | **<0.001** | 0.431 | 0.971 |
| Control-High | 1.571 | 0.407 | -2.201 | 0.133 |
